# Mechanical forces instruct division plane orientation of cambium stem cells during radial growth in *Arabidopsis thaliana*

**DOI:** 10.1101/2024.05.23.595483

**Authors:** Mathias Höfler, Xiaomin Liu, Thomas Greb, Karen Alim

## Abstract

Precise regulation of cell division is central for the formation of complex multicellular organisms and a hallmark of stem cell activity. In plants, due to the absence of cell migration, correct placement of newly produced cell walls during cell division is of eminent importance for generating functional tissues and organs. In particular, during radial growth of plant shoots and roots, concerted cell divisions in the cambium are essential to produce adjacent xylem and phloem tissues in a strictly bidirectional manner. While several intercellular signaling cascades have been identified to instruct tissue organization during radial growth, a role of mechanical forces in guiding cambium stem cell activity has been frequently proposed but, so far, not been functionally investigated on the cellular level. Here, we coupled anatomical analyses with a cell-based vertex model to analyze the role of mechanical stress in radial plant growth at the cell and tissue scale. Simulations based on segmented cellular outlines of radially growing *Arabidopsis* hypocotyls revealed a distinct stress pattern with circumferential stresses in cambium stem cells which coincided with the orientation of cortical microtubules. Integrating stress patterns as a cue instructing cell division orientation was sufficient for the emergence of typical cambium-derived cell files and agreed with experimental results for stress-related tissue organization in confining mechanical environments. Our work thus underlines the significance of mechanical forces in tissue organization through self-emerging stress patterns during the growth of plant organs.

## 1 Introduction

Cambium-driven radial growth is a fundamental growth process in many plants increasing the circumference of their shoots and roots by the production of vascular tissues. In the form of wood, vascular tissue production contributes by a large proportion to the world’s production of biomass ^1^, hence, also acts as an important long-term sink of atmospheric CO2^2^. Radial growth generally is driven by cambium stem cells (CSCs). CSCs reside in a cellular environment hosting very rigid tissues while generating xylem and phloem cells in a bi-directional manner ^3^. Xylem cells are usually produced inwards and ensure long-distance water transport as well as provide mechanical stability by their well-developed secondary cell walls. Phloem cells are produced outwards and play an important role in transporting sugars and other organic molecules ^3–5^. Cambium-derived cells often display very organized alignments in radial orientation (Figure 1A). This organization is based on a strong preference for tangential (periclinal) divisions of CSCs for which mechanical forces have been proposed to be instructive ^6^. As one of many possible alternative explanations, concentration gradients of peptides secreted by phloem cells have been suggested to serve as signals controlling the oriention of CSC divisions ^7^.

**Figure 1.**
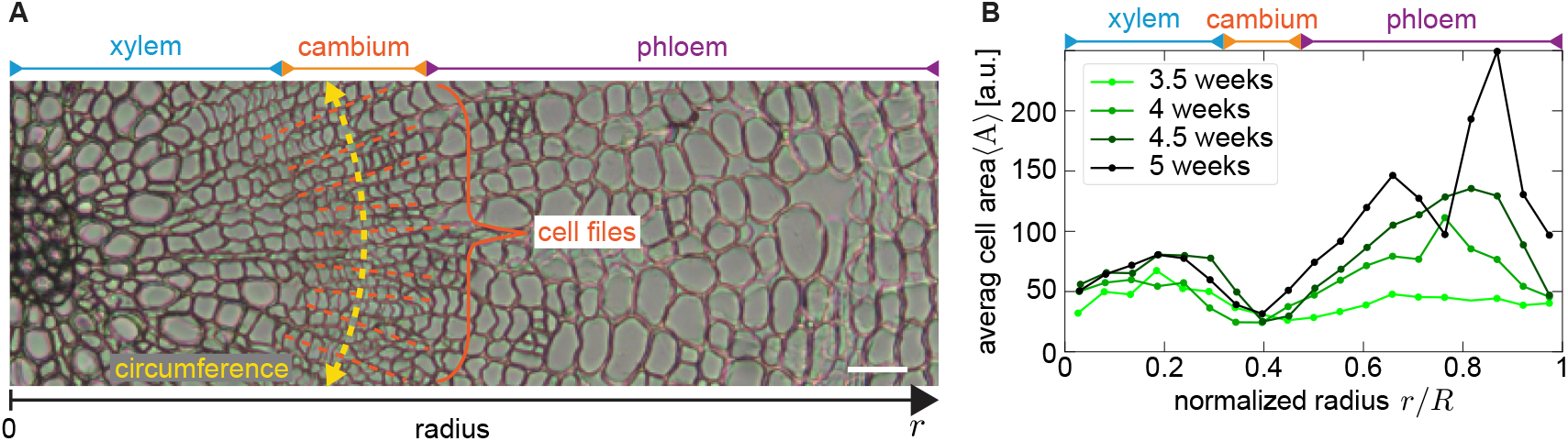
Cambium tissue is organized into cell-files and cell areas in the hypocotyl show a bi-modal distribution. (**A**) Cross section of an *Arabidopsis* hypocotyl. Cambium-derived cells are arranged in cell files (yellow dashed lines) showing a high level of organization, compared to xylem and phloem tissue. Scalebar 10 µm. (**B**) Binned average cell area distribution (bin size 0.05) as a function of the normalized tissue radius *r/R* with respect to the tissue center shows a bi-modal distribution at all ages. Underlying data from ^11^.

The hypocotyl of *Arabidopsis thaliana* is the organ connecting the shoot and the root system and serving as a versatile model for investigating radial plant growth, including morphometric analyses and *in silico* simulations of cambium-based organ expansion ^8–12^. These simulations have partly assumed inter-tissue forces generated by developing xylem cells instructing CSCs’ division orientation ^10^. However, the pattern of mechanical forces in radially growing plant organs have not been revealed yet, let alone their impact on dividing CSCs in a natural context.

Various division rules were proposed for plant cells in the past. The most prominent of those rules is that cells divide along the plane that minimizes the surface area of the new cell plate (Errera’s rule) deriving from the observation that cells often behave like surface-minimizing soap bubbles ^13–17^. Still, cells also often deviate from this ‘shortest axis’ rule and purely geometrical rules usually do not accurately capture the dynamics of division plate placement ^6,18–20^. While auxin signaling was shown to promote deviations from Errera’s rule ^21^, there is, however, strong evidence that plant cells respond to mechanical stresses on a local or even tissue level, possibly impacting cell divisions ^6,16,18–20,22–35^.

### Box 1.

**Cell-based vertex model description**

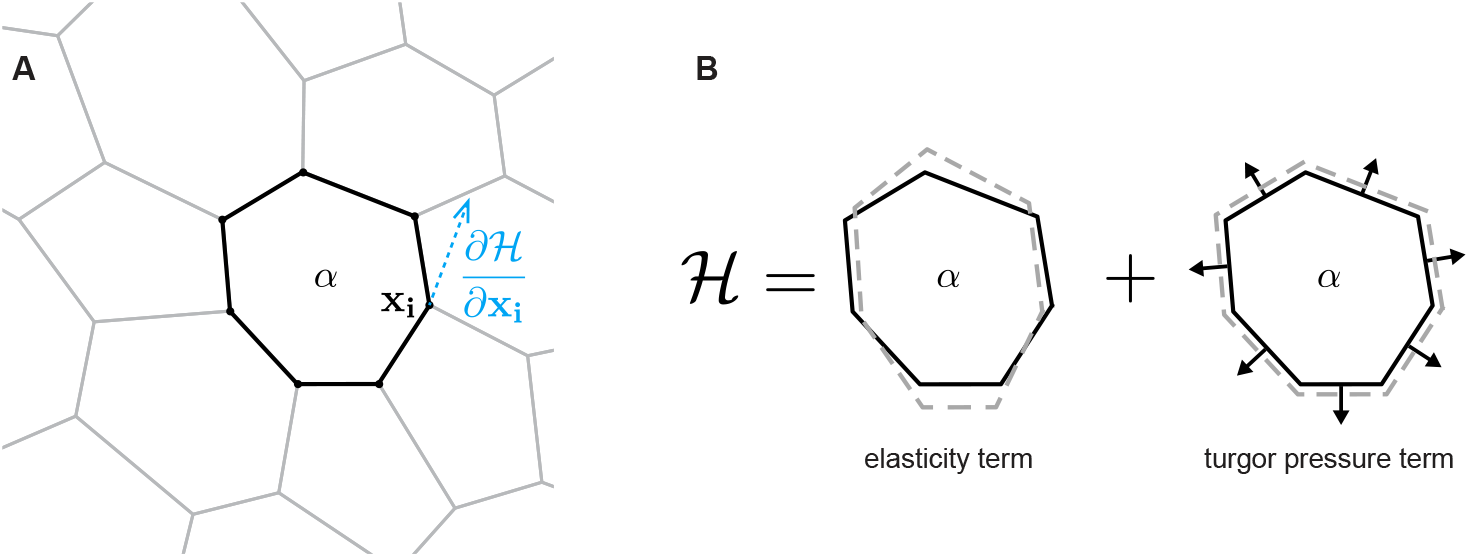

**Vertex model theory:**

The tissue is described by a cell-based vertex model in two dimensions. Cells are represented as a graph of connected vertices, defining encircled polygons (**A**). Polygons have an area *A* and are numbered by *α* = 1 … *N*. The cell mechanical properties are represented by the energy function ℋ, consisting of a term for the elastic energy of cells and pressure within cells (**B**)

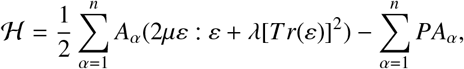

with strain *ε*, Lamé constants *µ* and *λ* and turgor pressure *P*. : denotes the Frobenius norm of *ε*. We follow a quasi-static approach in order to find stable tissue configurations that satisfy force balance such that the sum of forces *F*_*i*_ at each vertex position *x*_*i*_ vanish, thereby locally minimizing the energyℋ (*x*_*i*_). The first sum of the energy function represents the elastic energy of the tissue taking into account the Lamé parameters *µ* and *λ* and the strain *ε* acting on each cell *α*, which is derived from the mismatch between the observed cell shape and the cell shape at minimized energy. The second sum is a volume term, assuming a constant cell height normalized to one, and represents the turgor pressure of the cells.

Plant cell shapes depend largely on internal turgor pressure and the mechanical properties of surrounding cell walls ^36^. Mechanical forces not only emerge from individual cellular properties, though, but especially from the tissue- and organ-wide arrangement and expansion of cells due to the tight connection of cells via their walls ^22,37,38^. Furthermore, various cell structures align with the direction of maximum tensile stress, most notably cortical microtubules (CMTs) which orient parallelly to the stress vector ^17,32,39–43^. This implies that tension caused by mechanical changes in neighboring cells, for example through ablation or other mechanical cues, manifests by a reorientation of CMTs ^44^. As a consequence, the orientation of cell division planes may thereby be affected ^17,32^.

In this work we investigated if mechanical stresses, arising during radial growth, are instructive in orchestrating coordinated tissue growth and arrangement. To this end we started with a quantitative analysis of radial growth of the *Arabidopsis* hypocotyl. By performing two-dimensional simulations of cross sections starting from segmented cell outlines within a vertex model, we found a distinct pattern of circumferential stress in CSCs and radial stress in the xylem and phloem. Our simulations revealed that stress-oriented cell divisions explain the emergence of ordered tissue structures during radial growth and, thus, that organ-wide stress patterns are prominent candidates for decisive positional cues during cambium-dependent tissue production. This conclusion was suported by disrupting tissue stress patterns through applying exogenous mechanical stress which led to the same disorganized tissues pattern in both simulations and experiments. Based on these observations, we propose that cell divisions oriented along mechanical stress vectors are central for organized radial growth of plant organs.

## 2 Results

### Cell-based vertex model captures radial hypocotyl growth

To assess the disparity in tissue type growth for deducing organ-wide mechnical stress patterns, we took advantage of previously published morphometric analysis of hypocotyl cross sections ^11^. As is shown in Figure 1B, we found both xylem and phloem cells to be much larger in surface area compared to cambium cells. Thus, mapping out the average cell area ⟨*A*⟩ versus the normalized radius *r/R* from the tissue center, with R being the total radius of the tissue, we found the area distribution to be bimodal. It was noticeable that the average xylem cell area increased only very slowly over time, while phloem cells grew much more within the same time interval. CSCs, on the other hand, kept on average the same size throughout tissue development.

To initialize mechano-simulations by a cell-based vertex model, we first segmented an image of a cross section (Figure 2A) from an Arabidopsis hypocotyl using the PlantSeg/MorphographX software packages ^45^ in order to obtain a realistic tissue mesh. As a compromise between accuracy and numerical robustness, from this segmentation we generated a Voronoi mesh (Figure 2B), using the center of mass for each cell in the segmentation. This resulted in a simplified tissue structure that retained, up to minor deviations, the quantitative characteristics of the original tissue, like the average cell area as a function of the radius (long/short axis orientation, Figure S2A) and the cell anisotropy (Figure S2C,D), i.e. the orientation of the short axis of each cell. In our model, we represented each cell by a polygon, defined by a series of vertices, connected by straight lines, i.e. edges (Box 1A). As contributions to the tissue energy we took into account the turgor pressure and the elastic energy of each cell (Box 1B). The elastic energy comprised the strain of each cell, which was proportional to the mismatch between the observed, i.e. deformed, shape of the cell *M*_*c*_ and the undeformed cell shape *M*_0_ (Figure 2C and Eq.2). The undeformed cell shape hereby evolved in time by applying exponential growth to the system (Eq.4). We followed a quasi-static approach such that in each simulation step, an optimization was performed in order to find a local minimum of the tissue energy, *H* (Eq.5). This corresponded to a stable configuration in which the forces on each vertex was balanced, resulting in a vanishing net force. For cell divisions, we defined division planes through the center of the dividing cell where the direction of division was determined depending on the division rule applied. The strain *ε* acting on the mother cell before the division was conserved such that both daughter cells combined experienced the same strain as the mother cell had been subject to before the division. This was achieved by matching the undeformed shape matrices of daughter cells to the undeformed shape matrix of the mother cell (Eq.10). In this way, daughter cells both inherited proportionally the amount of strain, depending on the ratio of their surface areas (Figure 2E). It was only after the division event, when a new local energy minimum was sought, that the tissue adapted to the newly introduced degrees of freedom and the polygonal network adapted accordingly to the mechanical forces.

**Figure 2.**
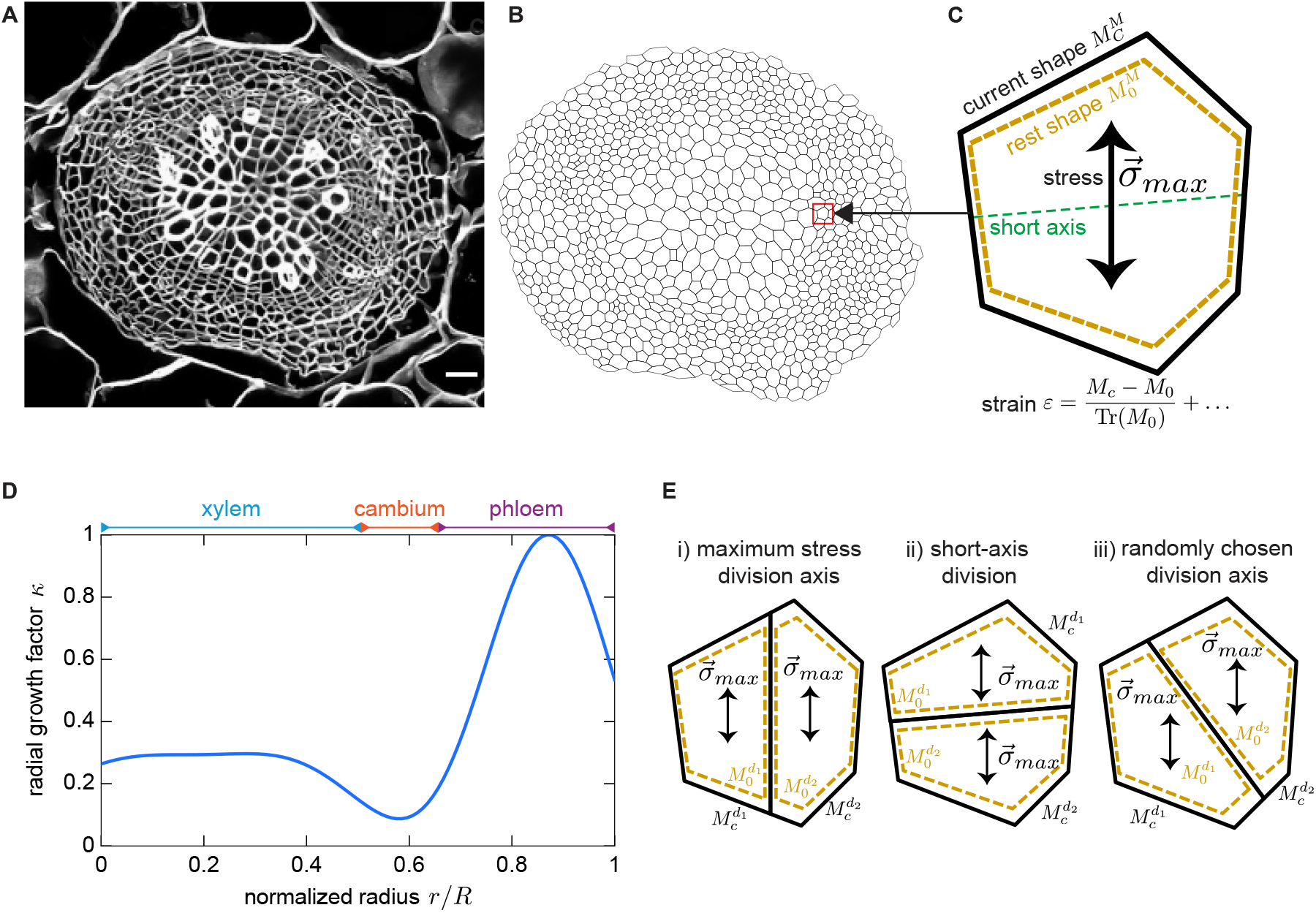
*In silico* recreation of segmented tissue and observed cell growth rate form basis for tissue level simulations, including cell divisions. (**A**) Cross section of a two-week old *A. thaliana* hypocotyl that served as simulation template. Scalebar 15 µm (**B**) Voronoi tessellation as simulation input, obtained from segmentation of A. (**C**) Description of individual cell geometries by means of shape matrices. (**D**) Radial growth function used to modulate individual cell growth as a function of radius. (**E**) Schematic of maximum stress, short-axis and random division rules for division plane placement, investigated in simulations.

Because our simulations intended to study the role of mechanical stresses in driving cell patterns in cambium-drived tissues, we focused on the *in silico* emergence of tissue structures during the early growth phase of the hypocotyl. A major challenge in this undertaking was the lack of many desired input parameters for the simulations such as the mechanical properties of different cell or tissue types and their dynamics during cell division and tissue development. We therefore used available data ^11^ in order to indirectly extract this information. All cells were mechanically treated equally in the simulations and we did not distinguish between different types. Differences in mechanical properties of the cells were taken into account indirectly by applying a radius-dependent growth factor *κ*(*r/R*) (Figure 2D) where *r* was the Euclidean distance to the tissue center of mass and *R* the radius of the tissue. This captured the observed evolution of the radial distribution of cell areas ^11^. The cell area distribution in both younger and older hypocotyls was considered bimodal ^46^ (Figure 1B, S2A, S3A) and differed in the average cell area of xylem and phloem tissue over time. Morphometric data ^11^ moreover suggested a massive cell growth in phloem tissue, implying the growth rate in the phloem to be larger than in the xylem. For this reason, the xylem makes up a comparably larger proportion in younger hypocotyls while, during aging, the phloem area massively increases. Therefore we used a growth function *κ*(*r/R*) which replicated this transition. For determining this function, we started to fit the sum of two Gaussians to the area data of our Voronoi mesh, obtained from segmentation. These took the form 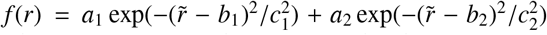 where 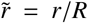 was the normalized radius. We combined the fit of the Voronoi mesh at 2 weeks of age (Figure S2A) with the experimental data at later stages (Figure 1B) to compute a growth rate that evolved the area distribution of the young sample’s Voronoi mesh such that it eventually matched the experimental data after growth. Since we were assuming a Gaussian distribution we could determine the slopes of amplitudes *A*_*i*_ and means *µ*_*i*_ for each distribution of the experimental data (Figure S3B,C) over time and considered this the rate of change of the respective quantity within half a week of growth. This slope was added to the parameters that described the Gaussian fit of the area distribution of the Voronoi mesh (Figure S3D), therefore corresponding to the expected area distribution at the sample age of 2.5 weeks. Additionally we adjusted the fit’s variance such that the x-position of the maxima and the local minimum of the original distribution of the Voronoi mesh were retained. This was necessary as increasing the amplitudes would have otherwise artificially shift the extreme values due to the summation of both Gaussians. Taking the difference of this distribution, corresponding to a 2.5 weeks old sample, and the distribution at the age of 2 weeks, we obtained a radial growth function *κ*(*r/R*) (Figure 2D, S3D) which we used to modulate individual cell growth, depending on the radial position of the cell center.

### Circular stress-patterns emerge from radially varying growth rates

Employing our model, we first looked into the mechanical stress patterns emerging dynamically within the tissue without cell divisions. The experimental observation of a bi-modal distribution of cell areas at all hypocotyl stages indicated xylem and especially phloem cells to be expanding faster than cambium cells. To implement the larger expansion rates, we increased for the simulations’ initial condition the undeformed shape matrices *M*_0_ for both xylem and phloem isotropically by 5% and 20%, respectively. Apart from this increase, we did not impose further initial conditions or any stress patterns. Instead, the initial stresses were determined from the energy optimization during the first simulation step. To quantify the emerging stresses for the subsequent analyses, we always used the tissue center as a reference point. Moreover, we averaged over 100 independent runs where individual cell growth underlied multiplicative Gaussian noise beyond the radial growth function *κ*(*r/R*). This ensured stochastic growth of each cell. The stress angle (Figure 3A), i.e. the angle of the direction of maximum stress, we defined with respect to the periclinal direction of the tissue, therefore the normal to the radius vector. For each individual cell, we computed the stress angle over the course of the simulation. In detail, we started by growing the tissue, obtained from the Voronoi tesselation of the segmented data, with a timestep of Δ*t* = 0.01. We then applied our radius dependent growth factor *κ*(*r/R*) in every simulation step and continuously monitored both the average stress angle 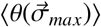 and the area distribution *A* as a function of the normalized radius *r/R* (Figure S2E,F). During growth of the initial mesh we observed the appearance of a distinct stress pattern, establishing already after a few simulation steps and resulting in a clear concentric domains (Figure 3B). Our quantitative simulation analysis (Figure S2E) showed that already shortly after the beginning of the simulation, at *t* = 1.0, there was a distinct difference between the average stress angles 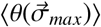 along the hypocotyl radius. At small radii up to about *r/R*≈ 0.6 stresses were largely oriented in periclinal direction whereas for larger radii, there was a clear transition towards the anticlinal direction. This pattern became more prominent over time (Figure S2E), also indicated by more pronounced transitions between periclinal and anticlinal stress along the radius of the tissue. For very small radii *r/R <* 0.2, due to the very small number of cells, we could not finally conclude the stress orientation, however the data suggested a rather radial orientation as well. Following our differential growth function, xylem and phloem cells grew both faster than cambium cells (Figure S2F), with phloem cells growing much faster than xylem cells, as desired from considering the experimental data (Figure 1B). Based on these observations we concluded that applying our growth function *κ*(*r/R*) on the initial conditions, derived from experimental data, circumferential stress in the cambium and radial stress in the xylem and phloem emerges.

**Figure 3.**
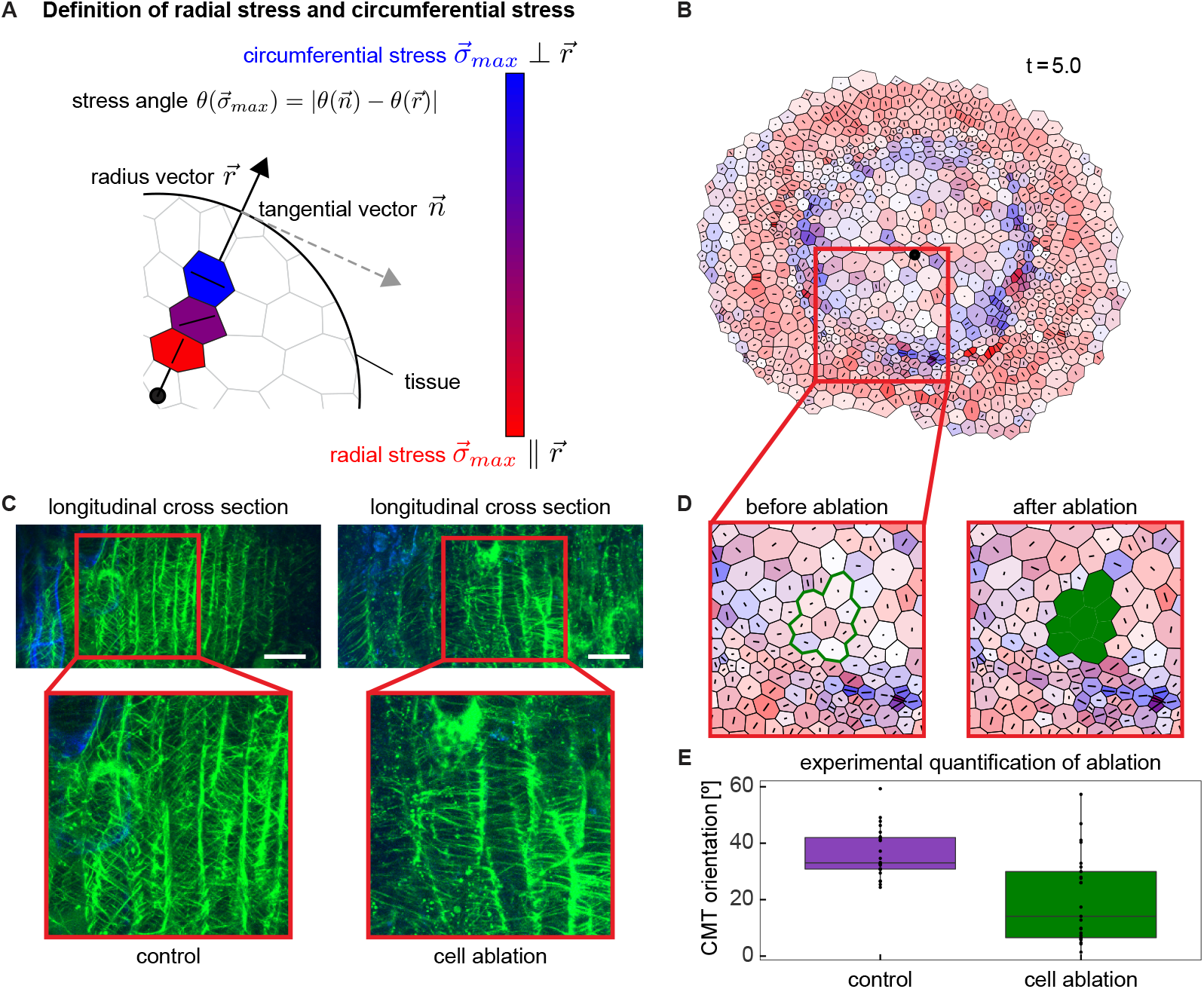
Simulations show a distinct stress pattern with circumferential stresses in the cambium and radial stresses in the xylem and phloem that respond also to mechanical perturbations. (**A**) Definition of the radial stress and circumferential stress, as well as the stress angle 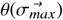. Measures are always with respect to the tissue center. The color bar captures the variation in stress between the extreme cases of the vector of maximum stress 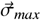 having either only a component in radial 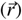 or circumferential 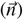 direction. (**B**) Distinct circumferential stress pattern after growing the initial Voronoi mesh (Figure 2B) at *t* = 5 (500 iterations). Tissue center marked by black dot. (**C**) Longitudinal cross sections of the hypocotyl with lignin in xylem vessels visualized by auto-fluorescence (blue) and CMTs visualized by*pWOX4::mVenus-MBD*-derived signals (green)l. Scalebar 25 µm. In the control (left) microtubules are spirtally winded around cells, after xylem removal (right), they are horizontally oriented, indicating a mechanical reinforcement towards the cavities. (**D**) *In silico* ablation of xylem tissue. After ablation (right), both xylem and cambium cells surrounding the ablation site, are subject to higher stress overall and stresses re-orient parallel to the area of ablation. (**E**) Quantiative experimental analysis of CMT orientation angle w.r.t. horizontal axis of control and after ablation.

To challenge the relevance of the observed stress pattern, we next compared the effect of *in silico* removal of the xylem tissue in the simulation with the removal of the xylem tissue *in planta. In silico*, we simulate the cell ablation by setting the stiffness of cells to zero and reducing the turgor pressure to a residual ten percent ^40^. Doing so, we observed an overall increase of stress around the ablation site (Figure 3D, right) even in cells at a few cell layer distance from the direct neighbors of the ablation site. There, stresses of surrounding cells reoriented around the wound such that the maxmimum stress was parallel to the ablated tissue surface. Xylem removal also led to increased circumferential stress in the cambium after removing the xylem tissue (Figure 3D). To see whether our simulations faithfully predicted stress dynamics *in planta*, we experimentally visualized CMTs in cambium cells in longitudinal cross sections of the hypocotyl. To this end, we used a line expressing the fluorescent mVENUS-MICROTUBULE BINDING DOMAIN (mVenus-MBD) ^47^ fusion protein under the control of the cambium-specific *WUSCHEL RELATED HOMEOBOX 4* (*WOX4*) promoter ^3^. After manual removal of the xylem tissue and a one-day recovery phase, we analyzed CMT organization in CSCs by live-cell imaging. Compared with non-disturbed controls, where CMTs showed a prominent spiral twisting (Figure 3C, left, E, left), CMTs in ablated samples were arranged more horizontally (Figure 3C, right, E, right). These observations let us conclude that, as predicted *in silico*, the xylem tissue applies pressure to CSCs preventing their radial stretching. We also concluded that our simulations, although abstracting the hypocotyl tissue as a two dimensional cell mesh, correctly predicted enhanced circumferential stress specifically in CSCs.

### Division rules are a key factor in tissue organization and division plane orientation

Having found support for the conclusion that our simulations faithfully captured tissue-wide stresses in the hypocotyl, we next employed our pre-simulated tissue mesh (*t* = 5.0, Figure 3B) to investigate the impact of cell divisions on tissue morphogenesis. To this end, we compared three different rules for cell division orientation: **i)** cells divide along the direction of maximum stress, **ii)** cells divide with random orientation, and **iii)** cells divide along their shortest axis (cf. Cell division). For all cases, we imposed the following constraints: Only CSCs were allowed to divide, but among CSCs, cells were picked randomly for initiating cell division. As CSCs, we considered cells in a radial range of *r*_*min*_*/R* ± 0.075*r/R*, where *r*_*min*_ was the minimum of the area distribution, corresponding to the center of the cambium domain. Due to the faster phloem growth as compared to the cambium, the local minimum of the area distribution was shifted towards the organ centre (Figure S2F, S4B,D,F). To ensure that only cambium cells divided during tissue growth, we tracked the minimum of the area distribution to dynamically keep the same reference for cell divisions throughout the simulation. In addition, cells only divided if they exceeded a certain size threshold, which was half of the average cambium cell size at the beginning of the simulation (*t* = 0). This rule was based on experimental data (Figure 1B) showing that cambium cell size remains on average largely constant, suggesting an upper size limit. Also, there was always only one division event per step and divisions were carried out only every 20 integration steps (Δ*t* = 0.2). These rules ensured tissue growth in addition to the divisions and allowed the tissue to integrate effects of one division before possibly undergoing another one in their nearest neighbors.

Simulations following the three different rules generated different cellular patterns. When cells divided in parallel to the maximum stress, division planes oriented largely periclinally at the beginning of each simulated division following the observed stress pattern (Figure 3B). When dividing randomly, resulting division planes generated angles wih a uniform distribution. For dividing along the shortest cell axis, division planes were determined by taking the eigenvector of the smallest eigenvalue of the shape matrix *M* of the mother cell (cf. Cell division). In both our Voronoi meshes and experimental data, application of this rule resulted largely in anticlinal divisions (Figure S2B,C,D).

As we were especially interested in tissue topology and cell division orientation, we compared these features generated under the different division rule regimes (Figure 4) with experimental data. Interestingly, stress patterns in our simulations remained intact independently from the chosen division rule (Figure 4C,E,G and S4A,B,C) indicating that not CSC behaviour but the faster growth of xylem and phloem tissue compared to the cambium (Figure 1B) was the driving factor for the emergence of these patterns. For our comparisons, division angles were defined as angles between the division planes and the direction 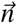 normal to the radius vector.

**Figure 4.**
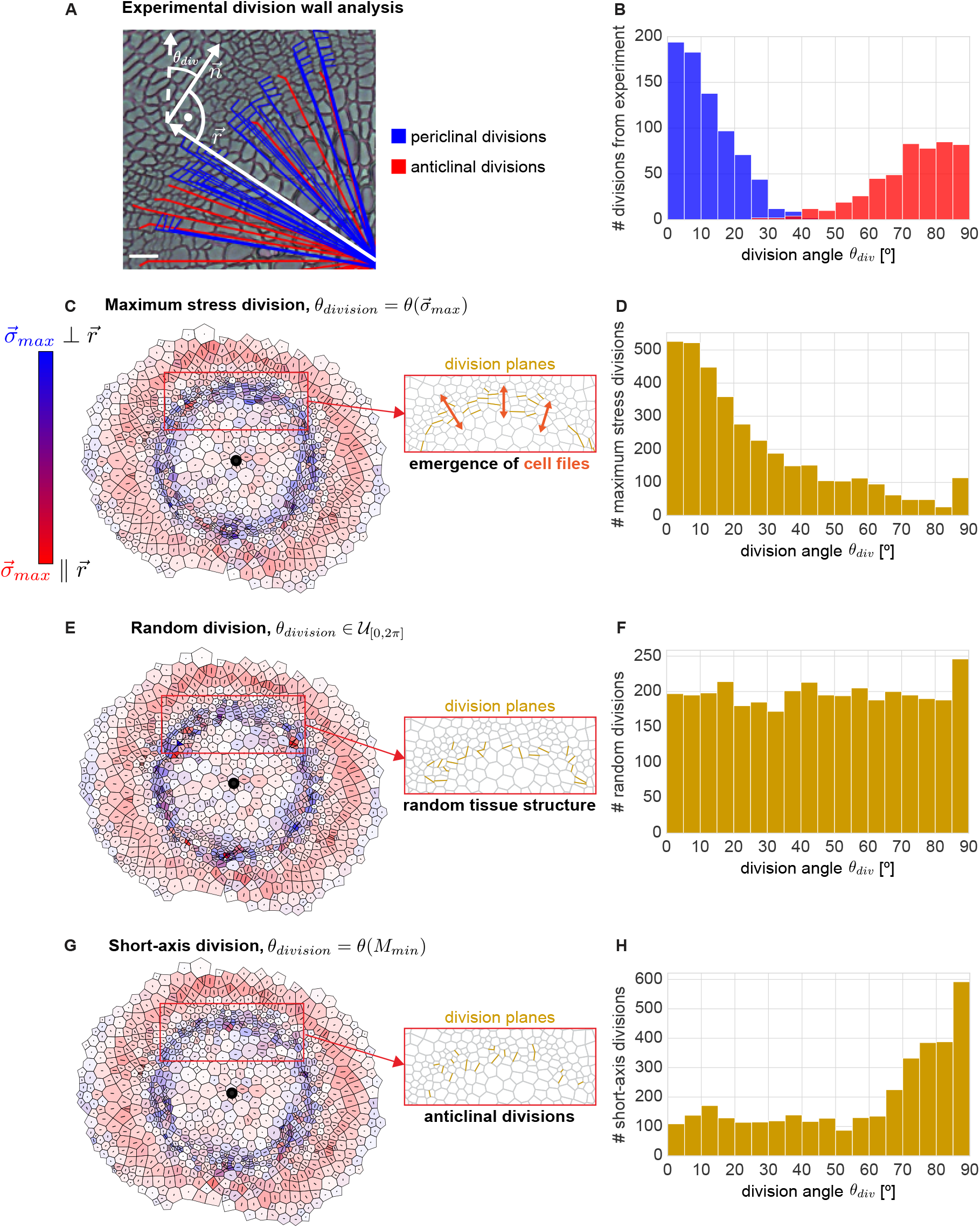
Experimentally measured peri- and anticlinal divisions correlate to maximum stress and short-axis divisions. (**A**) Depiction of experimental acquisition of division plane angles w.r.t. the normal of the radius vector 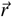. Scalebar 10 µm (**B**) Histogram of the division plane angles *θ*_*div*_ as measured in 1A(top) with periclinal (blue) and anticlinal (red) divisions. (**C**,**E**,**G**) Representative examples of tissues after 50 cell divisions with the maximum stress, random, and short-axis division rule, respectively. The insets show a magnification of parts of the cambium with division planes indicated in mustard-yellow. Cell files only emerge in the case of maximum stress division (C,D). (**D**,**F**,**H**) Histograms of division angles *θ*_*div*_ in direction of maximum stress, random direction and short axis direction respectively, with the angle being w.r.t. the periclinal direction, i.e. normal to the radius vector. Displayed is an average over 70 independent simulation runs.

Those planes were measured from one edge of an individual cambium cell to the center of mass of whole cross sections (Figure 4A). Quantitative analysis of division angles in plants (Figure 4B) showed that CSC divisions typically occured either with a periclinal (0° w.r.t. radius vector normal 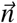) or anticlinal orientation (90° w.r.t the radius vector normal 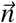). Division plane orientations in the simulations were analyzed in the same way (cf. Figure 3A). When doing so, we found striking differences between the different division rule regimes with regard to tissue topologies and distributions of division angles. (Figure 4D,F,H). For divisions along the maximum stress, we mostly obtained periclinal (Figure 4D) divisions and it was only for this division rule that we observed the highly regular cell arrangement with emerging cell files as observed in plants. The large variation in the angles, reflected by the broadness of the histogram distribution, arised from local perturbations in the stress pattern and shape anisotropies. These may have led to high strains and stresses in individual cells, imposed by or imposing onto neighboring cells, which did not necessarily align with the overall stress pattern of the tissue. When letting cells divide randomly, we observed an almost uniform distribution of division angles and a rather irregular arrangement of cells (Figure 4E). For short-axis divisions, anticlinal divisions were clearly the most prominent category and there was no clear topological pattern emerging (Figure 4G). While random divisions showed, as expected, a uniform distribution, short-axis divisions lead to an angular distribution that was complementary to the result of maximum stress divisions. This was a result of the short axis of cells pointing almost exclusively in radial direction which was largely perpendicular to the direction of maximum stress (cf. Figure S2C,D).

In conclusion, while all division rules left the stress pattern intact, only divisions along the direction of maximum stress led to the emergence of radial cell files in cambium-derived tissues as observed in plants. Therefore, our results demonstrate that divisions along the direction of maximum stress are strong candidates for directing cell divisions in CSCs and highlight the significance of mechanical forces towards tissue morphogenesis. As an interesting consideration, a combination of the maximum stress and short-axis division rules may serve as an explaination for the distributions of division plane orientations that were obserevd in plants. In that sense, mechanical stress may be the factor guiding cell file emergence, while the geometric short-axis rule is a candidate for guiding anticlinal divisions increasing the CSC pool in circumferential orientation.

### Mechano-perturbations lead to similar tissue organization changes *in silico* and *in planta*

To probe the significance of the alignment of CSC divisions along the direction of maximum stress, we investigated CSC divisions and vascular tissue organization in hypocotyls under exogenous mechanical stress. Therefore, we constrained hypocotyl expansion by positioning a clip applying pressure from two opposite sides of the hypocotyl after four weeks of non-perturbed growth (Figure 5A, Figure S1A). As the organ continued to expand, growth was thus constrained in two directions, resulting in regions with apparent anatomical differences (Figure 5A, white dashed lines). The regions of sections wich were positioned adjacent to the clip showed a very regular tissue structure, similar to the structure observed under regular growth conditions (cf. Figure 1A) and harbored typical radial cell files (Figure 5C,a). In comparison, in the regions which were positioned perpendicular to the clip (Figure 5C,b) tussie anatomy was very chaotic and no cell files were present. The more regular tissue anatomy next to the clip was not due to a general suppression of CSC divisions under the physical confinement, though, because the activity of *pHTR13::pHTR13-mCherry* reporter indicating indicating S-phase and early G1 phase during cell divisions and part of the PlaCCI (plant cell cycle indicator) ^48^ was comparably active under perturbed and non-perturbed conditions (Figure S1B,C). Hence, the high regularity of division plane orientation in CSCs adjacent to the clip did not simply reflect the situation before the start of the treatment. We further checked cambium organisation under clip treatment using combined *pPXY* and *pSMXL5* reporters visualizing cambium organization ^3^ and found that the typical arrangemnt of cambium domains was highly disturbed in regions positioned perpendicular to the clip (Figure S1E,F). Since CMT orientation is parallel to the maximum stress direction and directly influences cell division orientation ^49^, we next checked CMT organization in CSCs in the different regions of perturbed hypocotyls. These analyes showed that CMTs were more isotropic in CSCs perpendicular to the clip possibly refelcting an altered physical environment in the regions (Figure S5).

**Figure 5.**
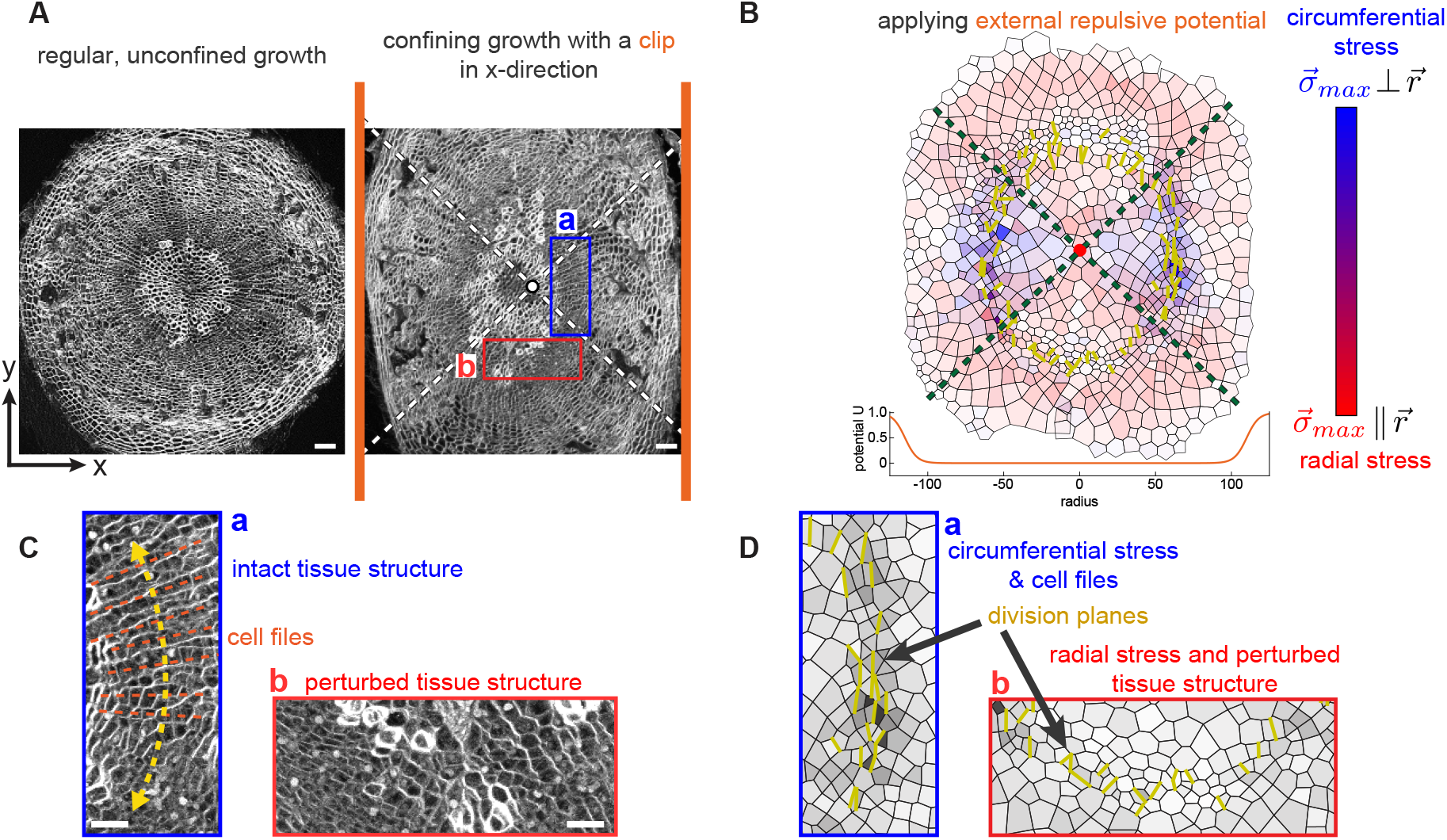
External mechanical stress instructs cambium organization by perturbation of division wall orientation. (**A**) Cross sections of hypocotyls under regular, unconfined growth condition (left) and under exogenous mechanical stress through confinement by clip treatment in x-direction (right). White lines correspond to quarters of tissue where the tissue structure remains well-organized (a, and directly opposite tissue quarter) or where the tissue structure is perturbed, respectively (b, and directly opposite). Scalebar: 50 µm (**B**) Implementation of the clip treatment in simulations. Growth is confined in x-direction by adding an external potential of logistic shape, acting as a soft wall on both sides of the simulated tissue. Shown is the tissue from Figure 3B after growth within the external potential (*t* = 20) and 75 cell divisions following the maximum stress rule. (**C**) Magnifications of cross section (A, right) with intact tissue structure including cell files (a) and perturbed tissue structure (b). Scalebars: 25 µm (**D**) Magnifications of two sections of the simulated tissue. Yellow lines represent the division planes in the cambium. Division planes are oriented almost exclusively in y-direction as a result of the applied external potential, overriding the original stress pattern of the tissue in b and the quarter directly opposite of b (cf. Figure 3B). This is accompanied by an intact tissue structure and the occurrence of cell files in D,a and a perturbed structure in D,b.

In simulations, we used the same initial mesh as for the previously studied cell divisions (Figure 3B) and applied a confining external potential *U*(*r/R*) in the shape of a logistic function (Eqn. 11). This confined growth in x-direction analogously to the *in planta* experiments, allowing growth in y-direction only (Figure 5B). When applying again the maximum stress cell division rule for 75 divisions under these conditions, we obserevd largely disrupted stress patterns in the regions positioned perpendicular to the confinement. After few rounds of simulation, stresses started to realign towards the y-direction, disrupting the concentric pattern of radial stress. As in the *in planta* experiments, we divided the tissue in four quarters (Figure 5B, green dashed lines). CSCs within the two quarters being directly subjected to the external constraint effectively remained under circumferential stress (Figure 5D,a) while CSCs located in the two quarters perpendicular to the exogenous stress were under radial stress (Figure 5D,b). For the two quarters maintaining circumferential stress, the simulation revealed almost unchanged stress patterns compared to our previous simulations manifested by the emergence of radial cell files due to periclinal cell divisions. On the contrary, for the two quarters in which CSCs experienced radial stress (Figure 5D,b) anticlinal cell divisions emerged preventing cell file formation very much in accordance with the *in planta* results. Therefore, we concluded that our simuations faithfully reported stress patterns in radially growing hypocotyls and that there is a functional connection between tissue organization and organ-wide mechanical stress patterns which emerge through a combination of locally defined cell production and expansion rates.

## 3 Discussion

Radial growth makes a vital contribution to the production of biomass on earth, however an in-depth understanding of the regulation of cell divisions and the influence of different mechanical environments is yet unknown. In this study, we combined *in planta* experiments and *in silico* simulations to understand how mechanical stress regulates cell division plane orientation in CSCs and general tissue organization in radially growing plant organs. We developed a cell-based vertex model (Box 1) based on experimental recordings of hypocotyl growth (Figure 1B) in order to study mechanical stress patterns and their morphological effects on the emerging tissue structure. This endevor was achieved by taking into account experimental evidence to derive rules for cell growth rates and estimate initial conditions, including realistic meshes from segmentations. When applying these rules in simulations, we observed a rapidly emerging characteristic stress pattern with high circumferential stress in CSCs as a result of fast growing cells surrounding the stem cell tissue, possibly straining and thereby stressing CSCs accordingly. Regarding cell geometry, whereas in plants and in simulations the short-axis orientation of cambium cells was largely in anticlinal direction, the experimentally obtained division planes were mostly in periclinal direction. This discrepancy already hinted at a possible correlation between mechanical stress and cell division plane orientation, rather than divisions being guided by cell geometry alone. In simulations, we compared three different division orientation rules: maximum stress, random and short-axis. We observed that in all cases the stress pattern emerging during growth (Figure 3B) was the same and, thus, independent of the applied rules (Figure 4C-H). In contrast, only when applying the maximum stress rule we found tissue morphologies in agreement with *in planta* morphologies characterized by cell files emerging from mainly periclinal cell divisions. Periclinal divisions were also present in externally compressed hypocotyls. Specifically, compressed regions exhibited periclinal divisions in simulations and *in planta* experiments producing well-structured cambium-derived tissues with emerging cell files. In contrast, organ regions experiencing tensile stress exhibited anticlinal divisions and a comparatively distorted tissue morphology again in simulations and in plants. These findings strongly argue for a connection between tissue-wide mechanical stress and the orientations of cell divisions and a resulting instructive role of mechanical stress in tissue morphogenesis during radial growth.

Yet, it appears that geometry still affects division plane orientation as there was a high amount of anticlinal divisions observed in plants (Figure 4B) resembling short-axis-driven divisions observed in simulations (Figure 4H). This suggests that on the one hand, mechanical stress largely guides cell divisions and leads to highly ordered tissue structures through the emergence of radial cell files. On the other hand, short-axis-driven divisions may be part of the collection of divisions found in CSCs. Of note, the term ‘short-axis’ applies only when looking at the cambium in 2D cross sections. Because CSCs are very much elongated along the longitudinal organ axis, the shortest cell axis in fact lies perpendicular to this axis. Still, taking into account the aforementioned consideration, we suggest a combination of stress-based and geometry-based cell divisions in the cambium.

As CSCs are embedded deeply in growing organs, it is so far almost impossible to directly visualize them alive in their natural environment. Advanced experimental techniques like combined Raman and Brillouin microscopy ^50^ allow for the non-invasive acquisition of quantitative data about physical properties of tissues. Yet, due to limitations in light penetration it is also not possible for these techniques to collect physical data on CSCs in a natural non-perturbed context. Consequently, we believe that the simulation platform generated during this study will serve as a highly useful tool for investiagting mechanical stimuli in the context of radial plant growth. Getting a better understanding of tissue growth and division in the hypocotyl and other tissue types, especially the impact of mechanical stress towards these, will provide more insight into plant growth organization under specific environmental conditions. This could be a valuable insight for agriculture and forestry, helping to optimize growth in diverse ecological settings for example where mechanical stress might be a limiting factor, such as windy or rainy environments or the influence of gravity ^51^. With this work it furthermore becomes apparent that not only in the shoot apical meristem but also in other plant tissue, mechanical stresses, play a major role in tissue organization and the production of biomass via influencing cell divisions.

## STAR⋆METHODS

- Experimental procedures
  - Plant material and culture conditions
  - Vector construction
  - Histology
  - Confocal microscopy
  - CMT quantification in CSCs
  - Cell division angles quantification
  - Image and data analysis
  - Cell ablation
  - Clip treatment
- Theoretical methods and parameters
  - Vertex model
  - Stress and strain from shape matrix description
  - Framework and parameters
  - Cell division
  - External potential
- Data and code availability

## Supplemental information and data availability

Supplemental information, simulation code and experimental data, can be found online at: https://mediatum.ub.tum.de/1741765, using the login credentials.

Login: reviewer-access-04

Password: Lc95*pGEk/!tR%Kd8SpN

Reserved DOI: https://doi.org/10.14459/2024mp1741765.

## Acknowledgements

This work was supported by the German Research Foundation (DFG) through the FOR2581 grants GR2104/6, AL1429/4-1 and AL1429/3-1). We further want to thank João Ramos (Technical University of Munich, Germany) for plenty of fruitful discussions regarding his simulation model whose program code this work expanded on. We also thank Alexandra Zakieva (COS, Heidelberg University, Germany) for kick-starting this collaboration which allowed us to combine experimental data and simulations the way we could present it here.

## Author contributions

The concept for this work was created by all the authors, especially Karen Alim and Thomas Greb. *In planta* data were collected and analyzed by Xiaomin Liu and simulations were carried out and analyzed by Mathias Höfler. The manuscript was written by Mathias Höfler with support of Xiaomin Liu. All authors revised the manuscript.

## STAR⋆ METHODS

### Experimental methods

#### Plant material and culture conditions

*Arabidopsis thaliana* of Col-0 ecotype was used throughout this study. Arabidopsis seeds were surface sterilized with 70 % EtOH and 0.05 % Triton X-100 (Signa, T8787) and shaken for 1 or 2 minutes on a rotating wheel, the liquid was removed by a Pasteur pipet and 200 to 300 µl 100 % EtOH was added. Next, seeds were dried for one minute on a filter paper under sterile conditions. Sterilized seeds were sown onto sterile agar plates containing 1/2 Murashige and Skoog (MS) solid medium supplemented with 1 % sucrose, and stratified at 4°C for 2 days in the dark. Seedlings were transfered from plates to soil after 1 week in growth chambers under short day (SD) conditions (10 h light, 4 h dark, 22°C, 65 % humidity). 3-week-old seedlings were moved from SD to long day (LD) conditions (16 h light, 8 h darkness, 22°C, 50 % humidity).

#### Vector construction

For *pWOX4::mVenus-MBD* (*pXL12*), the following modules were combined in pGGZ001: WOX4pro(A), B-dummy(B), mVenus(C), MBD(D), WOX4 terminator(E) and hygromycinR(F) ^3,52–54^.

#### Histology

4-week-old hypocotyls embedded in JB4 resin (Polysciences, 00226-1) were used for histological analysis (Figure 1A, top) as previously described ^7^. 5 µm thick sections were made and visualized using bright field light. Images were captured by a Leica DM IRB microscope. The images shown in Figure 2A and Figure 5B; 5D are free-hand hypocotyl cross sections. For this, hypocotyls were embedded in 8 % Low Melting agarose (Sigma-Aldrich, A9414) in a small plastic weighing cup and cut with a razor blade (Wilkinson Sword). Sections were then put in a 2-well glass-bottom dish (ibidi, 80287) and stained with 0.1 % Direct Red 23 (Sigma-Aldrich, 212490) in Ca2+ Mg2+-free PBS for 5 min. After washing with water once, images were captured. For reporter lines (Figure S1B; Figure S2B; Figure S3B), free-hand hypocotyl cross or longitudinal sections were made as mentioned above, images were captured without staining.

#### Confocal microscopy

A Nikon A1 confocal microscope with a 25× water immersion objective lens (Nikon, Apo 25xW MP, 77220) was used to capture Figure 2A. The 561 nm laser was used to excite Direct Red 23. Hypocotyls of 2-week-old WT seedlings on plates under SD conditions were used. Images were collected at 2048× 2048 pixel resolution and 4× line average. A Leica Stellaris 8 confocal microscope with 20× air objective (HC PL APO CS2, 20X/0.75 DRY) or 63× water objective (HC PL APO CS2, 63× /1.2 WATER) and were used in all other images in this study. The Diode 405 nm and WLL lasers were used for excitation of fluorescent proteins or dyes. GFP was excited at 473 nm and detected at 480 nm to 529 nm. Citrine and mVenus were excited at 514 nm and detected at 558 nm to 615 nm. Autofluorescence of lignin in xylem vessels was visualized using a 405 nm laser and the emission was collected at 425 nm to 474 nm. Direct Red 23 was excited at 495 nm and detected at 558 nm to 615 nm. Image acquisition was performed at 1024×1024 pixel scanning field and 4× line accumulation.

#### CMT quantification in CSCs

Longitudinal sections of *pWOX4::mVenus-MBD* hypocotyls were used in this experiment. Pixel size in z dimension was set to 1 µm. CMTs organization of individual cambium cells was detected by choosing different stacks of images and making max projections in FIJI. Microtubules orientation and anisotropy of the selected CSC were taken from the ‘FibrilTool’ in FIJI as described before ^55^.

#### Cell division angles quantification

Images taken from JB4 resin sections were used to quantify cell division orientation in the cambium. FIJI was used to conduct images analysis by using the ‘angle tools’. For periclinal cell divisions, angles from the center of mass of cross sections to the tangent of the cell file (no less than 3 cells) direction were measured. Angles of anticlinal cell division were indicated by the angle between the center of mass of cross sections and the common cell wall of two cells which break up cell files. 15 independent hypocotyls cross sections were analysed and a final database of 751 or 498 division angles was produced as periclinal or anticlinal cell divisions, respectively.

#### Image and data analysis

The image in Figure 2A was processed as previously described to get the cell mesh ^11^. The PlantSeg parameters were set as follows: prediction patch size-1×128×128, stride-Accurate, segmentation algorithm-MutexWS, under-/over-segmentation factors-0.45, CNN, prediction threshold-0.06, Watershed seeds sigma-4.2, Watershed boundary sigma-4.2.

#### Cell ablation

The hypocotyl surface was exposed by cutting plants at the hypocotyl-root border. Next, the xylem tissue was removed using a needle. In controls, the hypocotyl surface was exposed but that xylem was left intact. Plants were then left recovering for one day in 1/2 MS liquid medium in growth chambers under LD conditions and free-hand sections using a razor blade (Wilkinson Sword) were produced. Those were immersed in water in a 2-well glass-bottom dish (ibidi, 80287).

#### Clip treatment

Home-made clips were applied to the upper parts of hypocotyls of 4-week-old seedlings grown in soil. After three days, clips were carefully removed and hypocotyls collected for histological analysis.

### Theoretical methods and parameters

#### Vertex model

Simulations employ a cell-based vertex model as introduced in ^56^ and furthermore applied in ^57–59^ that allows individual access to the mechanical properties of each cell within the tissue. A cell is represented as a pressurized container following the dynamics of a linearly elastic isotropic solid. Tissues used for the simulations are derived from experimental data, segmented with the software PlantSeg/MorphographX ^45^. The segmentation is used to obtain cell centers which themselves determine the Voronoi tesselations that serves as initial condition for simulations. Abstracting the recorded tissue by a Voronoi tesselation balances accuracy of tissue representation and numerical robustness as the individual shapes change, however, the experimentally recorded cell area distribution is maintained (Figure S2A).

#### Stress and strain from shape matrix description

Individual cells are described by shape matrices *M* and *M*_0_. *M*_0_ hereby represents the shape before deformation, i.e. the shape that minimizes the tissue energy *H* (Eq. 5) and *M* represents the shape after deformation defined by the second moment of area which in index notation is ^56,59^:

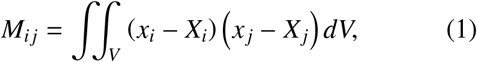

where *x*_*i*_ are the vertex coordinates of the cell and *X*_*i*_ the coordinates of the center of the cell. The formalism of employing the current shape *M* and the rest shape *M*_0_ allows to derive individual cell strain ^60^

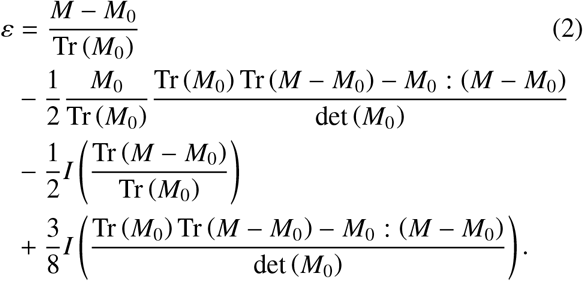

Strain determines stress from the constitutive relation

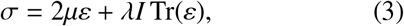

with Lamé parameters *µ* and *λ*. Cells grow exponentially by plastic deformation ^56,61,62^, driven from cell shape mismatch before and after deformation, *M*_0_ and *M*, respectively. Similarly to ^56^, *M*_0_ evolves as:

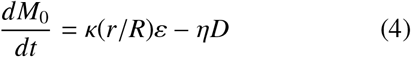

where the first term *κ*(*r/R*) represents strain-based growth with a cell-type specific growth factor here determined by a cell’s relative radius *r/R*. The second term models mechanical feedback mediated by CMT alignment with feedback constant *η* = 0.01 and the deviatoric stress tensor *D* which enhances growth in directions perpendicular to the maximum stress and reduces growth in the directions parallel to it as determined from the eigendecomposition of the stress tensor *σ*.

The total energy is described by ^56,60,63^

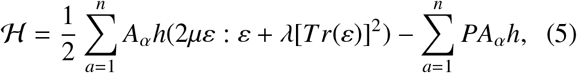

where the first sum is the elastic energy of the cells in the tissue and the second sum is the turgor pressure term, where *A*_0_ is the cell area in the undeformed state and *h* the cell height and equals one here since our model is strictly two-dimensional.

#### Framework and parameters

We use a C++ code, based on the quad-edge ^64^ data structure and its implementation as a C library ^65^. The energy function (Eq. 5) is minimized by using the C library NLOPT ^66^ with the NLOPT LD VAR1 algorithm, which is based on ^67^. ODEs in our model are solved with the GNU Scientific Library (GSL) ^68^, using the integrated Runge-Kutta-Fehlberg method (RKF45). Simulation parameters are determined as far as possible from the literature. Consulting ^69–71^, we apply a reference stiffness of *E*_0_ = 300 MPa and a turgor pressure of *P* = 0.5 MPA. The Poisson ratio we set to *ν* = 0.3.

Similarly to ^40^, for cell ablation we set the stiffness of ablated cells to *E*_*y*_ = 0 and set a residual turgor pressure to ten percent of the original value, i.e. *P* = 0.05 MPA.

#### Cell division

Cell division always occurs through the center of mass of the dividing cell. For random division, cells divide along a random direction chosen from a uniform distribution. For maximum stress divisions, we obtain the division plane orientation from the largest eigenvalue of the eigenvectors of the stress tensor *σ*. For short-axis division, we compute the eigenvector corresponding to the smallest eigenvalue of the cell’s observed shape matrix *M*. Upon cell division, we keep the strain of the mother cell conserved by daughter cells inheriting their *M*_0_ from the dividing mother cell. Since strain and stress directly depend on the current shape *M* and rest shape *M*_0_, we derive the new daughter cells’ rest shapes after division by transformation. As strains are very small, we transform between the current, deformed matrix *M* and the undistorted rest shape *M*_0_ with a single transformation Λ, employing the definition of infinitesimal strain from the displacement gradient. That is ^60^

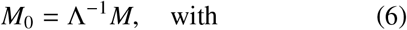

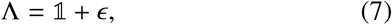

where Λ is determined from inversion of Λ by

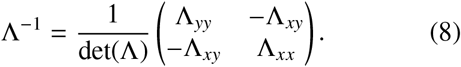

It then holds for all, the dividing mother cell and the resulting daughter cells, that

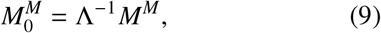

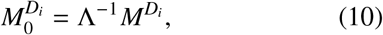

with the same transform Λ^−1^ which directly conserves the total strain in the system since it is just transferred to both daughter cells with the same transformation, thus also conserves total energy.

#### External potential

For the application of an external potential, we add a term *U* to the energy function (Eq. 5). For this we choose a logistic curve as it’s continuously differentiable and allows for a big slope to properly confine simulated tissue. This is defined as:

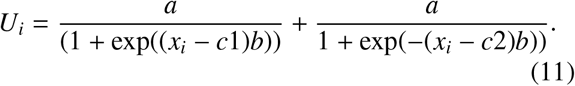

where *U*_*i*_ is the contribution to the energy by a vertex at position *x*_*i*_.

## Data and code availability

The data collected and simulation code used for this study is available online at: https://mediatum.ub.tum.de/1741765, using the login credentials.

Login: reviewer-access-04

Password: Lc95*pGEk/!tR%Kd8SpN In the future it will be openly available by the reserved DOI: https://doi.org/10.14459/2024mp1741765.

**Figure S1:**
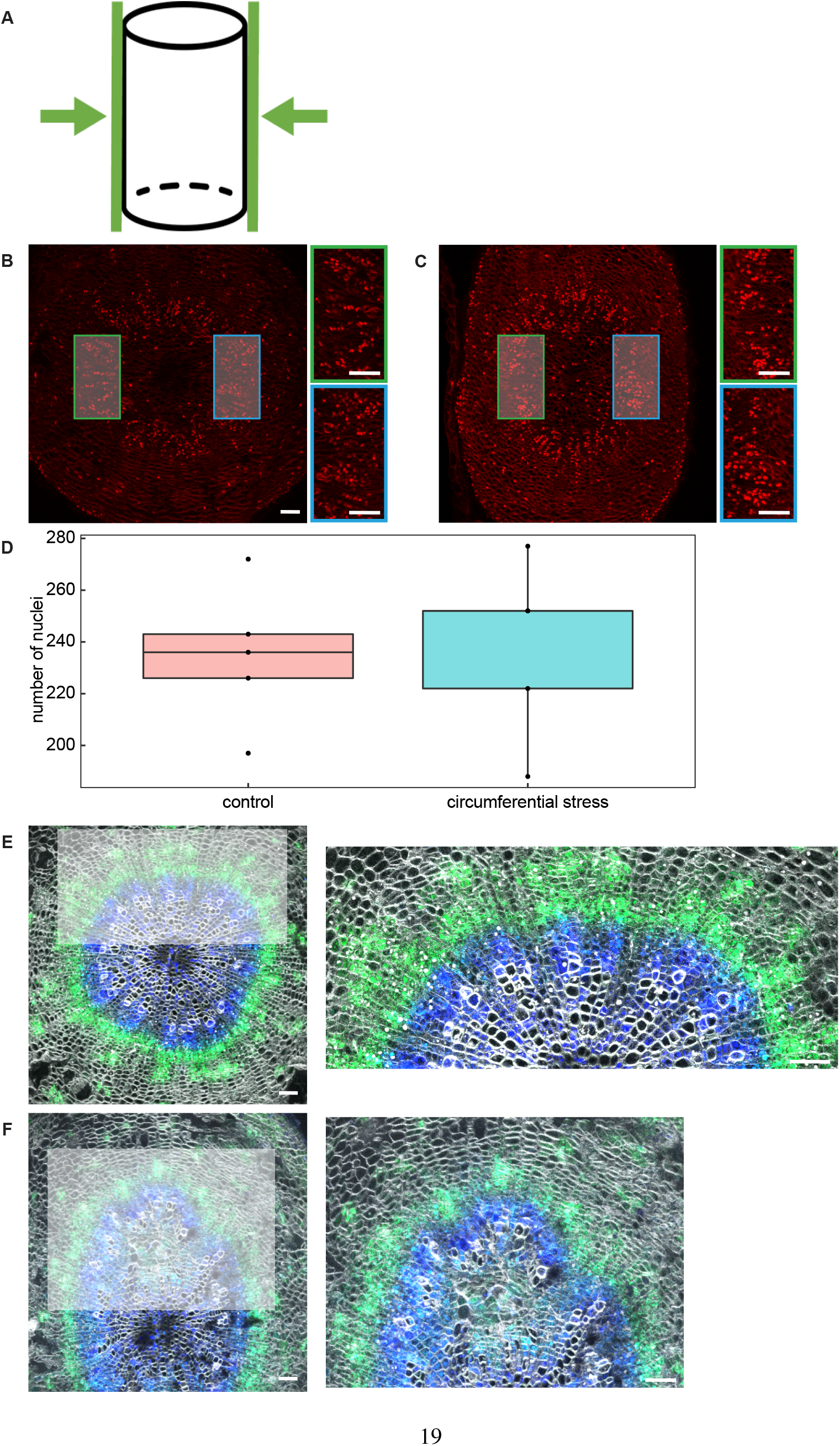
Effect of mechano-perturbation on cell division rates and cambium organization. (**A**) Schematic figure of exogenous mechanical stress treatment of the Arabidopsis hypocotyl. (**B**,**C**) Cross sections of the nuclei-localized *H3*.*1-mCherry* expression under control (B) or circumferential stress (C). (**D**) Quantification of the nuclei number (in the shadow area in B or C) of the *H3*.*1-mCherry* line under control or circumferential stress (n=5). (E,F) *pPXY::ER-mTurquoise, pSMXL5::ER-Venus* hypocotyl cross sections under control (**E**) or exogenous mechanical stress treatment (**F**). The images on the right are the magnifications of the shaded areas within the images on the left. Scalebar 50 µm

**Figure S2:**
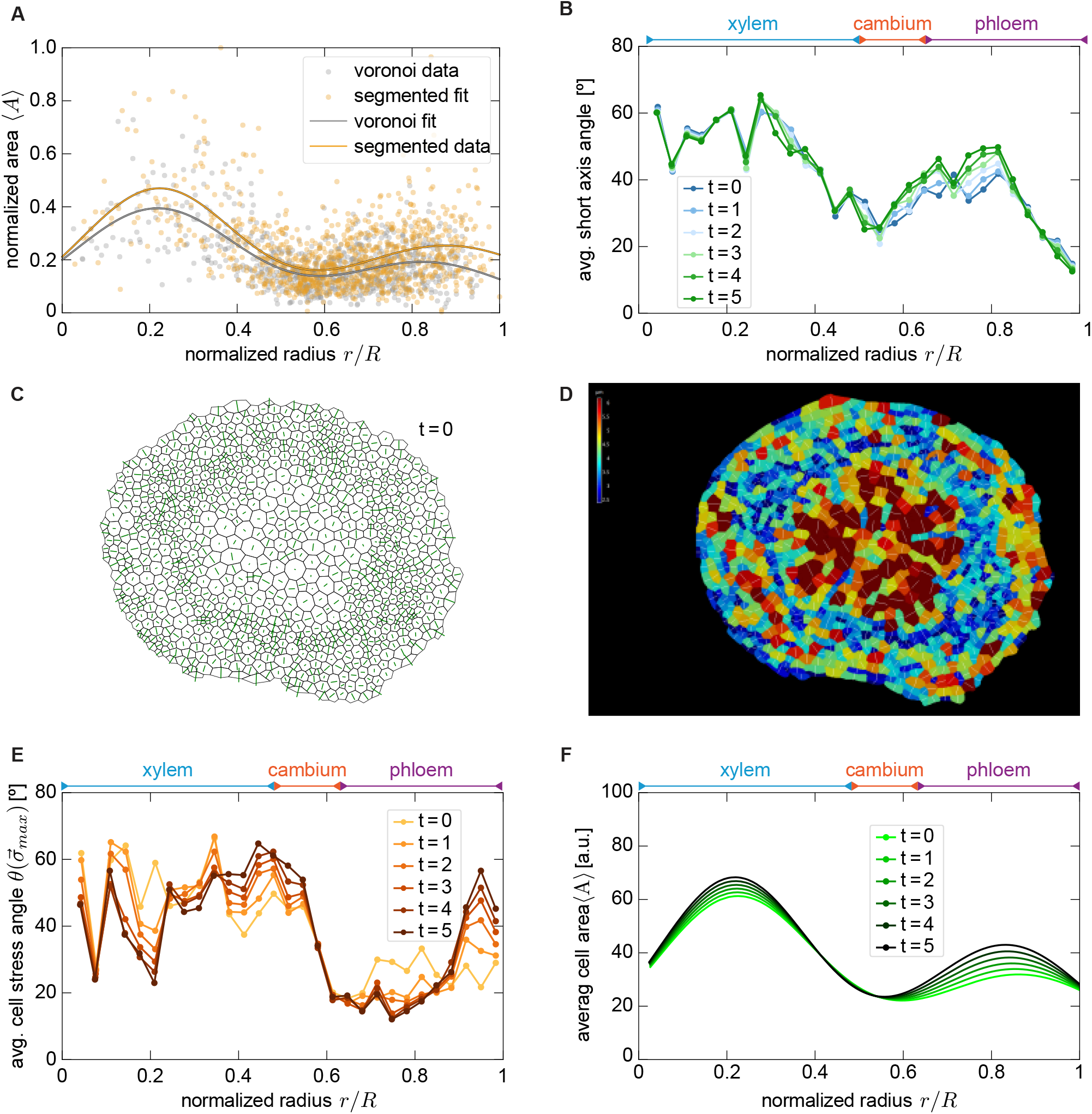
Simulation of the tissue as voronoi mesh retains the quantitative features of the original tissue and applying the derived growth function (Figure 2D) leads to sharp change in stress orientation in cambium and desired growth evolution of the simulated tissue. (**A**) Fit of the normalized individual cell areas *A* as a function of the normalized radius *r/R* for both the segmentation (yellow) and the derived Voronoi tesselation (grey) in comparison. (**B**) Angular orientation of short axis in simulation averaged and binned (bin size 0.033) as a function of the normalized radius *r/R*. (**C**) Voronoi mesh of simulation with short axes indicated (green lines). (**D**) Segmentation of 2 weeks old sample with indicated surface areas (colors) and short axes (white lines). (**E**) Evolution of averaged, binned (bin size 0.033) cell stresses. (**F**) Fit of individual cell areas as a function of the normalized radius *r/R*. For (B,E,F), we averaged over 100 independent runs.

**Figure S3:**
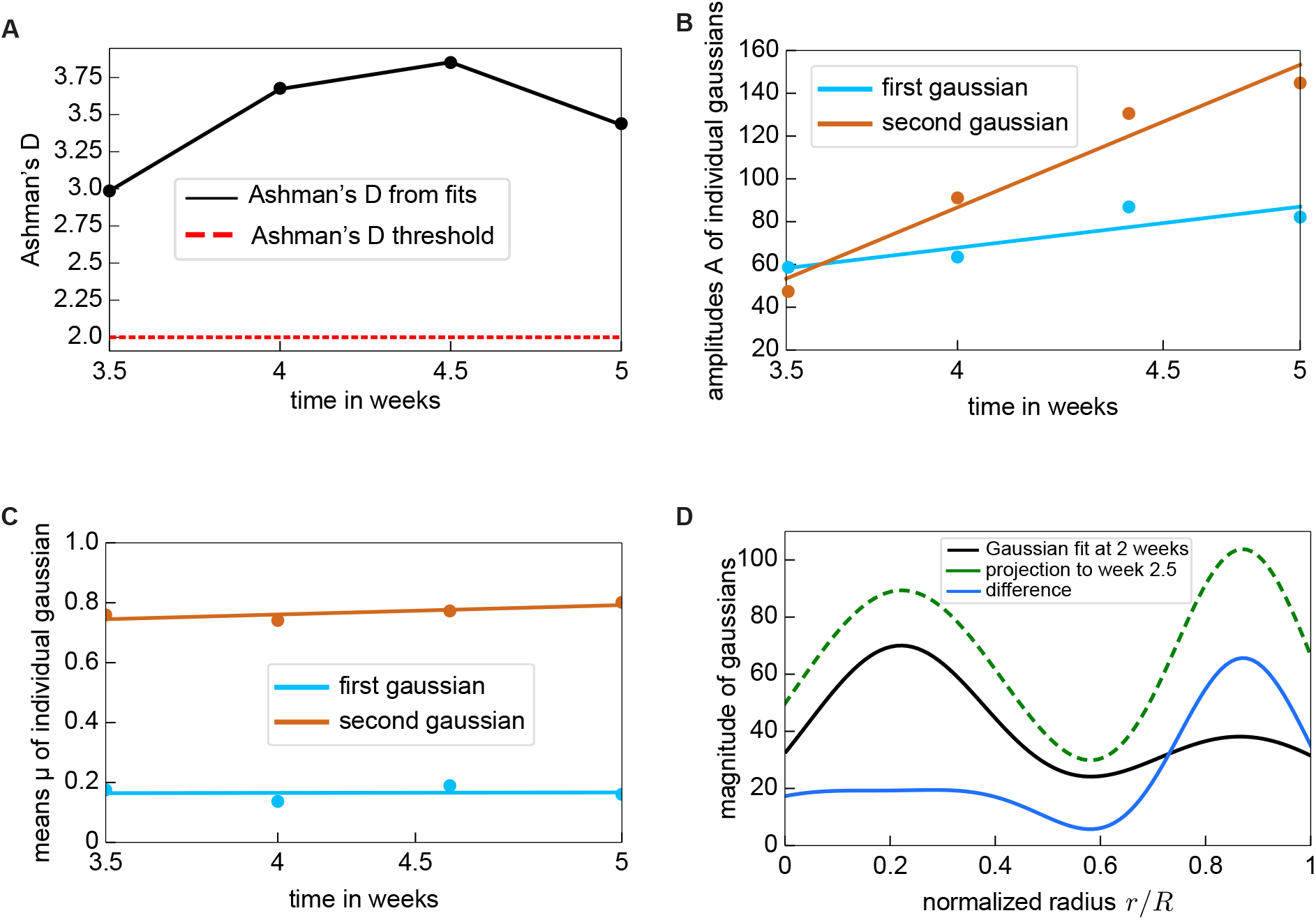
Bi-modal distribution of experimental data allows for derivation of radius-dependent growth factor. (**A**) Ashman’s D ^46^ for the experimental data in Figure 1B as a criterion that the average cell area distribution can be considered bi-modal. (**B**) Amplitudes of the Gaussian fits to the experimental data in Figure 1B with slopes *α* = 19.16 and *β* = 66.67. (**C**) Means of the Gaussian fits to the same data with a slope of *γ* = 0.0016 and *δ* = 0.031. (**D**) Gaussian fit (black) of the Voronoi mesh areas as in Figure S2A and the same fit function with adding the slopes from (A) and adjusting the variance to match the maxima and the local minimum of the Voronoi mesh areas. The difference of these curves results in the radial growth factor *κ*(*r/R*), we apply to individual cell growth in our simulations.

**Figure S4:**
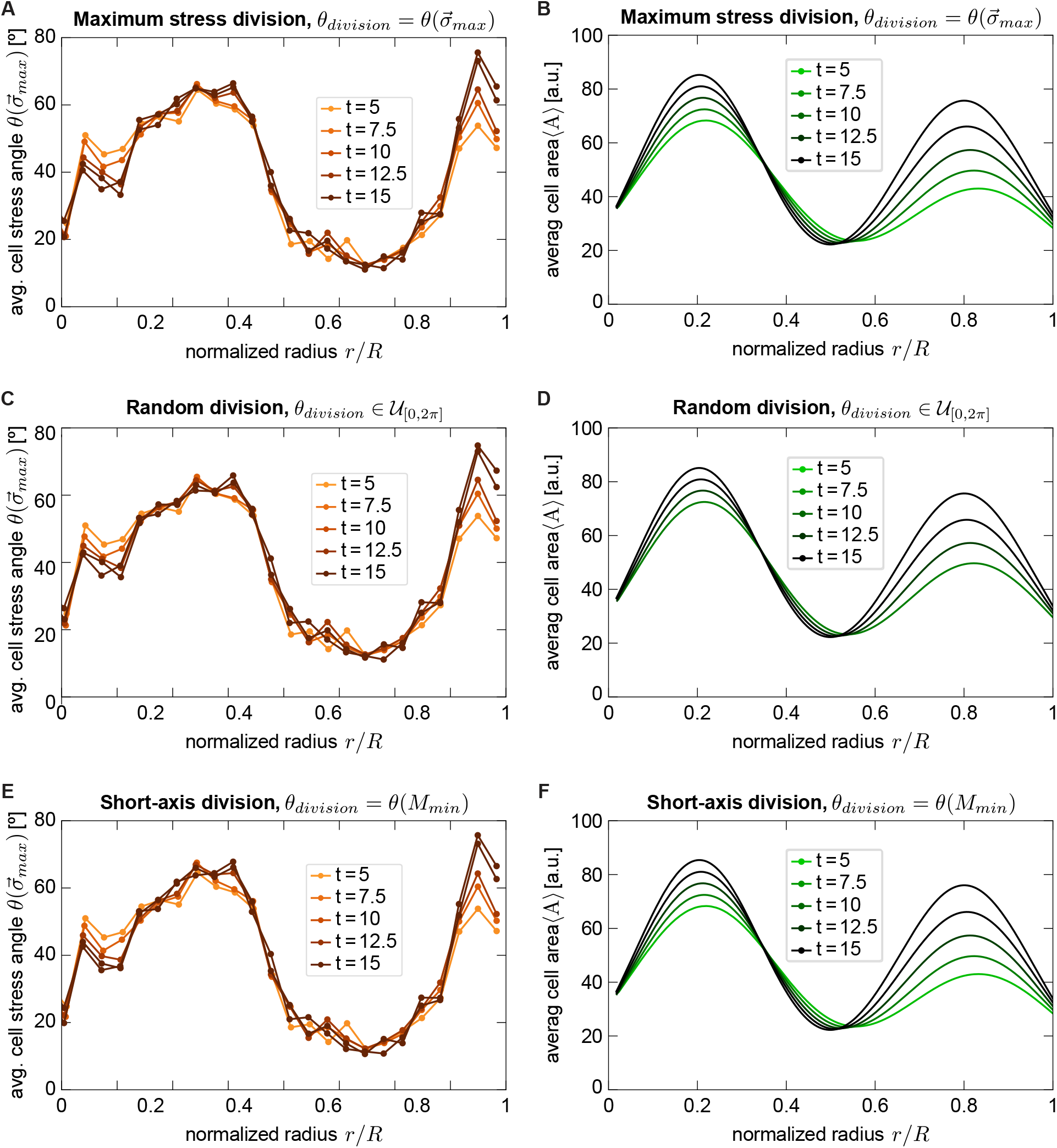
Different division rules do not largely change the stress pattern or cell area distribution. (**A**,**C**,**E**) Average stress angles w.r.t. radius vector 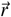 for random division, maximum stress division and short-axis division respectively at different time steps after the start of division (*t* = 5). (**B**,**D**,**F**) Fit of area distribution for all division rules at the same steps. For each plot we averaged over 70 independent runs.

**Figure S5:**
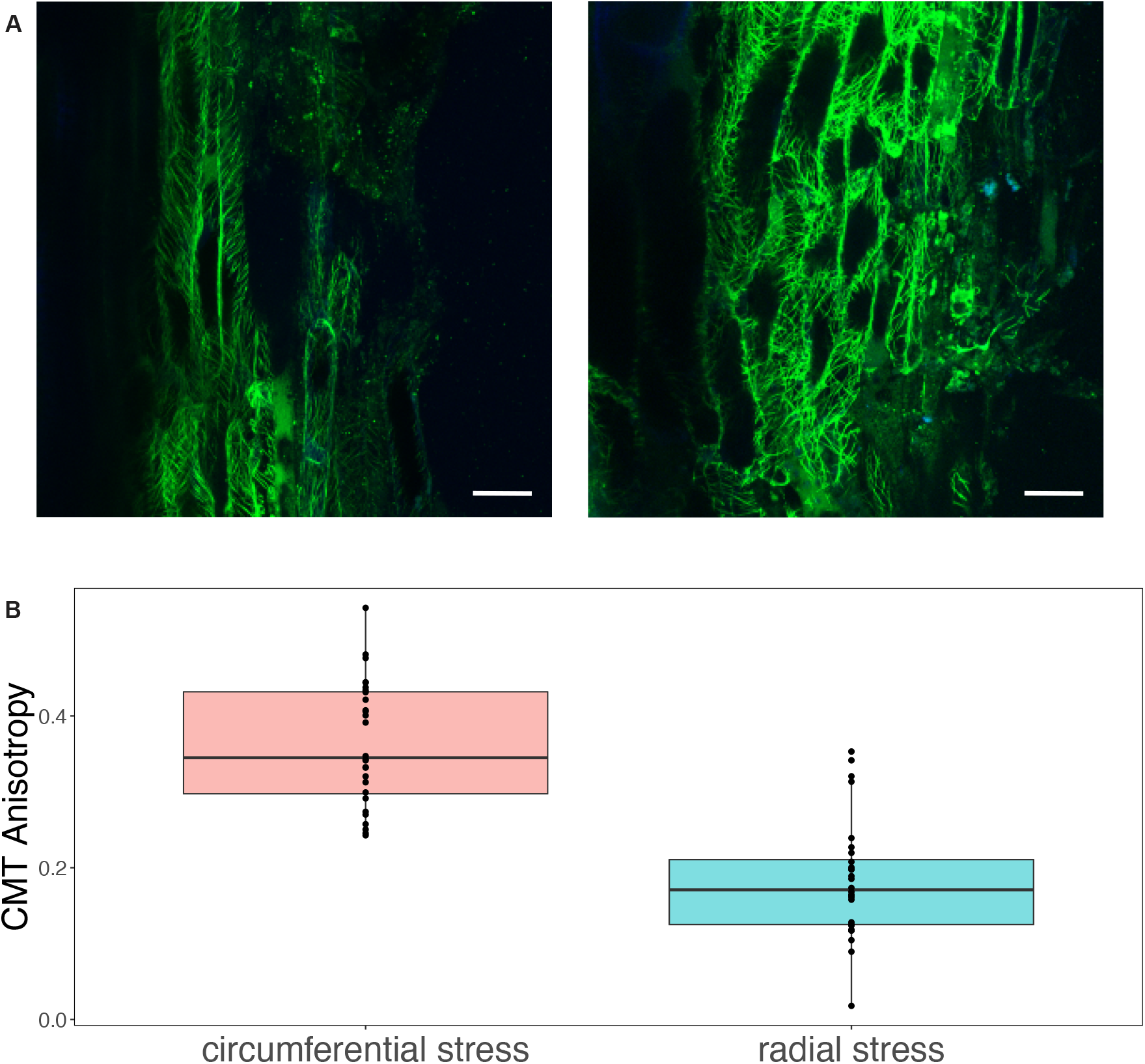
Microtubule orientaton in compressed versus non-compressed regions. (**A**) Longitudinal sections of hypocotyls under circumferential stress (left) or radial stress (right) using *pWOX4::mVenus-MBD*. Green, mVenus signal; (**B**) Quantification of CMT anisotropy in cambium stem cells under circumferential stress or radial stress. n=28 cells, P¡0.0001. Scalebar 20 µm.

